# The E2 glycoprotein holds key residues for Mayaro virus adaptation to the urban *Aedes aegypti* mosquito

**DOI:** 10.1101/2022.04.05.487100

**Authors:** Ferdinand Roesch, Chelsea Cereghino, Lucia Carrau, Alexandra Hardy, Helder Ribeiro-Filho, Annabelle Henrion Lacritick, Cassandra Koh, Jeffrey Marano, Tyler Bates, Pallavi Rai, Christina Chuong, Shamima Akter, Thomas Vallet, Hervé Blanc, Truitt Elliot, Anne M. Brown, Pawel Michalak, Tanya LeRoith, Jesse Bloom, Rafael Elias Marques, Maria-Carla Saleh, Marco Vignuzzi, James Weger-Lucarelli

**Affiliations:** Institut Pasteur, Viral Populations and Pathogenesis Unit, Centre National de la Recherche Scientifique UMR 3569, Paris, France; Institut Pasteur, Viruses and RNA Interference Unit, Centre National de la Recherche Scientifique UMR 3569, Paris, France; UMR 1282 ISP, INRAE Centre Val de Loire, Nouzilly, France; Department of Biomedical Sciences and Pathobiology, VA-MD Regional College of Veterinary Medicine, Virginia Tech, Blacksburg, Virginia, USA; Department of Microbiology, New York University Langone Medical Center, New York, United States of America; Brazilian Biosciences National Laboratory, Brazilian Center for Research in Energy and Materials (CNPEM), Campinas, SP, Brazil; Translational Biology, Medicine, and Health Graduate Program, Virginia Tech, Roanoke, Virginia, USA; Department of Bioinformatics and Computational Biology, School of Systems Biology, George Mason University, Fairfax, VA, USA; Program in Genetics, Bioinformatics, and Computational Biology (GBCB), Virginia Tech, Blacksburg, VA, USA; Research and Informatics, University Libraries, Virginia Tech, Blacksburg, VA, USA; Department of Biochemistry, Virginia Tech, Blacksburg, VA, USA; Edward Via College of Osteopathic Medicine, Monroe, LA, USA; Center for One Health Research, VA-MD Regional College of Veterinary Medicine, Blacksburg, VA, USA; Institute of Evolution, University of Haifa, Haifa, Israel; Basic Sciences Division and Computational Biology Program, Fred Hutchinson Cancer Research Center, Seattle, Washington, United States of America; Howard Hughes Medical Institute, Chevy Chase, Maryland, United States of America

**Author notes:** **Corresponding Author:** James Weger-Lucarelli.

## Abstract

Adaptation to mosquito vectors suited for transmission in urban settings is a major driver in the emergence of arboviruses. To better anticipate future emergence events, it is crucial to assess their potential to adapt to new vector hosts. In this work, we used two different experimental evolution approaches to study the adaptation process of an emerging alphavirus, Mayaro virus (MAYV), to *Aedes aegypti*, an urban mosquito vector of many arboviruses. We identified E2-T179N as a key mutation increasing MAYV replication in insect cells and enhancing transmission by live *Aedes aegypti*. In contrast, this mutation decreased viral replication and binding in human fibroblasts, a primary cellular target of MAYV in humans. We also showed that MAYV E2-T179N was attenuated *in vivo* in a mouse model. We then used structural and experimental analyses to show that MAYV E2-T179N bound less efficiently to human cells, though the decrease in replication or binding was not mediated through interaction with one of the host receptors, Mxra8. When this mutation was introduced in the closely related chikungunya virus, which has caused major outbreaks globally in the past two decades, we observed increased replication in both human and insect cells, suggesting E2 position 179 is an important determinant of alphavirus host-adaptation, although in a virus-specific manner. Collectively, these results indicate that adaptation at the T179 residue in MAYV E2 may result in increased vector competence – but coming at the cost of optimal replication in humans – and may represent a first step towards a future emergence event.

**Author Summary:** Mosquito-borne viruses must replicate in both mosquito and vertebrate hosts to be maintained in nature successfully. When viruses that are typically transmitted by forest dwelling mosquitoes enter urban environments due to deforestation or travel, they must adapt to urban mosquito vectors to transmit effectively. For mosquito-borne viruses, the need to also replicate in a vertebrate host like humans constrains this adaptation process. Towards understanding how the emerging alphavirus, Mayaro virus, might adapt to transmission by the urban mosquito vector, *Aedes aegypti*, we used natural evolution approaches to identify several viral mutations that impacted replication in both mosquito and vertebrate hosts. We show that a single mutation in the receptor binding protein increased transmission by *Aedes aegypti* but simultaneously reduced replication and disease in a mouse model, suggesting that this mutation alone is unlikely to be maintained in a natural transmission cycle between mosquitoes and humans. Understanding the adaptive potential of emerging viruses is critical to preventing future pandemics.

## Introduction

Arboviruses impose significant economic and public health costs worldwide. Zika virus (ZIKV), dengue virus (DENV), chikungunya virus (CHIKV) and West Nile virus (WNV) are now globally established and have caused significant outbreaks. In past decades, they mainly affected developing countries in tropical regions of the world and have been often neglected. Recently, local cases of ZIKV and CHIKV [1,2] have been detected in Europe and the United States, demonstrating that more temperate areas of the world are endangered by arboviruses, and that preventing arboviral emergence is one of the key public health challenges for years to come.

Mayaro virus (MAYV; Genus *Alphavirus*) was first isolated in Trinidad and Tobago in 1954 where it was associated with cases of mild febrile illness [3]. Since then, MAYV has caused sporadic outbreaks in several countries in South and Central America, including Brazil [4], Mexico [5], Peru [6], French Guiana [7], Bolivia [8], Ecuador [9] and Venezuela [10]. Imported cases have been described in multiple European countries including the Netherlands [11], France [12], Germany [13] and Switzerland [6]. Three genotypes have been described for MAYV: D (widely dispersed), L (limited) and N (new) [14]. However, recombinant strains of MAYV may also circulate in the Amazon basin [15]. Despite the increase in epidemiological data and surveillance in recent years, MAYV circulation is still likely under-estimated. Many factors contribute to this underreporting, including antibody cross-reactivity which complicates serological studies [6]; significant overlap in the areas of distribution of MAYV and CHIKV; and the important similarities between the clinical manifestations caused by MAYV and other arboviruses [16]. Indeed, like CHIKV and DENV, MAYV infection induces a febrile illness, with symptoms such as fever, rash, headaches, and nausea [17]. MAYV also induces arthralgia in most symptomatic patients [18] which sometimes lasts for several months [19] and may be caused by sustained production of pro-inflammatory cytokines [20,21]. In some rare cases, Mayaro fever is associated with more serious clinical outcomes, such as neurological complications, hemorrhagic manifestations or death [22]. Antibodies directed against MAYV have been observed in many animal species beyond humans, including primates, birds, and rodents [4,23] which may act as reservoir species.

MAYV is considered a serious candidate for viral emergence [14,24,25]. While most MAYV outbreaks so far have happened in rural areas in close proximity of tropical [26] or in indigenous communities [27], there is increasing concern that MAYV may escape this sylvatic cycle and become urbanized. Indeed, MAYV circulation has recently been observed in urban areas of Haiti [15]. In such areas, efficient transmission by *Aedes aegypti* would likely increase the risk of a sustained outbreak. Thus, the current epidemiological situation of MAYV is reminiscent of that of CHIKV before it emerged in 2006, when a single mutation in E1 allowed for better adaptation to *Aedes* albopictus, ultimately leading to the CHIKV outbreak in the Indian Ocean region [28].

While it is clear that MAYV can infect forest-dwelling *Haemagogus* mosquitoes in the context of its sylvatic cycle of MAYV [29], it is unclear whether MAYV can be transmitted in nature by mosquito vectors suited for urban transmission. Several laboratory studies have suggested that *Aedes aegypti* [30–32] and *Aedes albopictus* [33,34] can be readily infected by MAYV and may support viral transmission, albeit to different levels. One report also suggested that *Anopheles* mosquitoes may also participate in MAYV transmission [35]. While these studies suggest that MAYV has the potential to be transmitted by urban vectors, definitive evidence that this happens outside of laboratory settings is still lacking.

In this work, we sought to test the hypothesis that similar to CHIKV pre-emergence, MAYV is not yet fully adapted to *Aedes aegypti* transmission. We used two different *in vitro* evolution approaches to identify the genetic blocks that MAYV must overcome to better adapt to the anthropophilic vector *Aedes aegypti*. Deep mutational scanning and serial passaging approaches both identified T179N as a key mutation in the receptor binding protein, E2, resulting in increased MAYV transmission by *Aedes aegypti*. Interestingly, this mutation decreased viral replication in human cell lines and led to attenuation in mice, suggesting that adaptation of MAYV to this new vector may come at an evolutionary cost, which may explain why this mutation has not arisen yet in nature. Evolution at this residue may constitute an important first step towards MAYV emergence and should be monitored closely through increased surveillance in areas of sustained MAYV circulation.

## Results

### Natural evolution and deep mutational scanning enrich for similar high frequency mutations in MAYV

We identified mutations enabling MAYV adaptation to mammalian and insect hosts using two complementary approaches: naturally evolving virus through serial passaging and facilitating evolution through deep mutational scanning (DMS). Serial passaging was performed in BHK-21 (baby hamster kidney fibroblast), Aag2 (*Aedes aegypti* mosquito), U4.4 (*Aedes albopictus* mosquito), and 4a-3A (*Anopheles gambiae* mosquito) cells. We sequenced the virus using Illumina next-generation sequencing (NGS) after passage 1, 5 and 10 (**Fig. 1A**). All variants were below our threshold based on coverage depth and frequency in the passage 1 samples. As expected, most high frequency mutations were observed in the passage 10 samples, and the vast majority were found in the genes encoding for the envelope proteins, specifically E2 (**Fig. 1B-D**). We only present mutations in the envelope proteins since few mutations in other protein-coding sequences were identified. Of note, we observed a consensus change in 3/6 replicates from Aag2 cells (with frequencies ranging from 0.58 to 0.97) at nucleotide position 8918 (**Fig. 1D**), which resulted in a change within E2 at amino acid position 179 from a threonine (T) to either an isoleucine (I) or an asparagine (N). Mutations were also observed at this site for U4.4 cells (**Fig. 1C:** 2/6; range of 0.07-0.89), but not mammalian cells (**Fig. 1B**). Following passage in 4a-3A cells, we observed a mutation within E2 at amino acid position 232 resulting in a change from histidine (H) to proline (P) (**Fig. S1**). **Supplementary File 1** presents all the variants that passed our threshold, along with their frequencies and sequence coverage.

**Figure 1:**
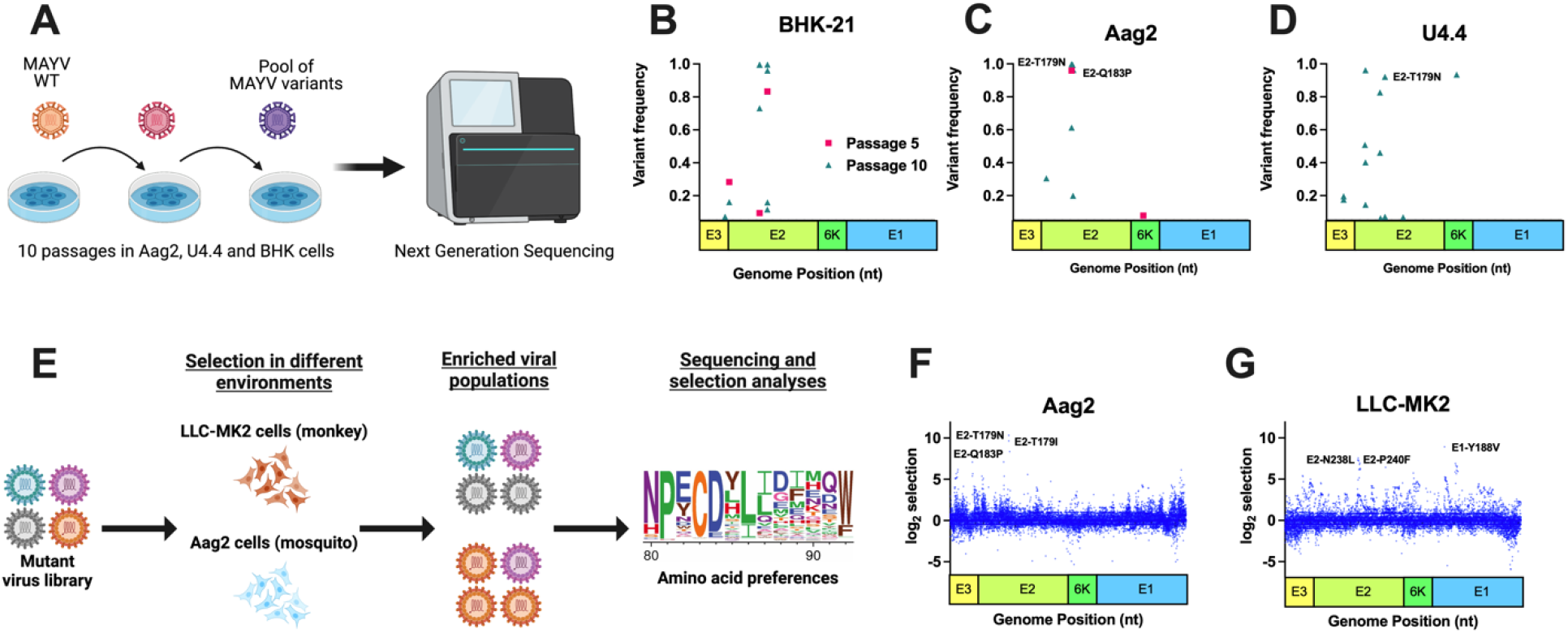
Natural evolution and deep mutational scanning (DMS) enrich for similar high frequency mutations in Mayaro virus (MAYV). Experimental evolution was performed using traditional serial passaging and DMS to identify molecular determinants of emergence. **A.** Traditional experimental evolution was performed by serially passaging MAYV at a MOI of 0.01 in BHK-21 cells (hamster) and two insect cell lines, Aag2 (*Aedes aegypti*) and U4.4 (*Aedes albopictus*). A total of ten passages were performed. **B-D.** Following one, five, and ten passages, the viral RNA was sequenced using Illumina NGS to identify potentially adaptive mutations. No high frequency variants were identified following passage one, and as such are not depicted here. **E.** The three MAYV DMS populations, along with WT MAYV, were used to perform three passages in Aag2 (**F**) and LLC-MK2 (**G**) cells. Following passage, the viral RNA was sequenced, and selection analyses were performed to identify enriched variants. The top three variants in each environment, based on selection strength, are presented for each graph.

MAYV DMS populations (**Fig. 1E**) were generated for all envelope genes (E3-E2-6K-E1; **Fig. S2A**) in 3 independent populations, as previously described for other viruses [36–38]. As expected, the initial DMS populations were highly diverse (**Fig. S2B**) as assessed by nucleotide diversity in the DMS and WT MAYV populations. We then passaged the DMS viruses in LLC-MK2 (monkey kidney epithelial) and Aag2 cells at a MOI of 0.01 for three passages. Virus titers were relatively stable throughout passaging in LLC-MK2 (**Fig. S2C**) and Aag2 (**Fig. S2D**). Following sequencing, we observed several amino acid changes that were enriched in Aag2- and LLC-MK2-passaged DMS populations (**Fig. 1F-G**). **Supplementary Files 2 and 3** present the full DMS selection outputs, including mutations enriched and selected against. Interestingly, the top three sites under selection (T179N, T179I, and Q183P) in Aag2 cells were in the E2 protein and were found as consensus changes in a least one of the replicates during natural evolution in Aag2 or U4.4 cells, though T179N and T179I were the most abundant in traditional passaging. The top sites under selection in LLC-MK2 were in E1 (Y118V) and E2 (P240F and N238L). We also passaged the DMS populations in mice and *Aedes aegypti* mosquitoes (**Fig. S2E-F**); however, only weak signals of selection were observed (**Fig. S3**). For the remaining studies, we will focus on mutations that were identified *in vitro*.

**Figure S1:**
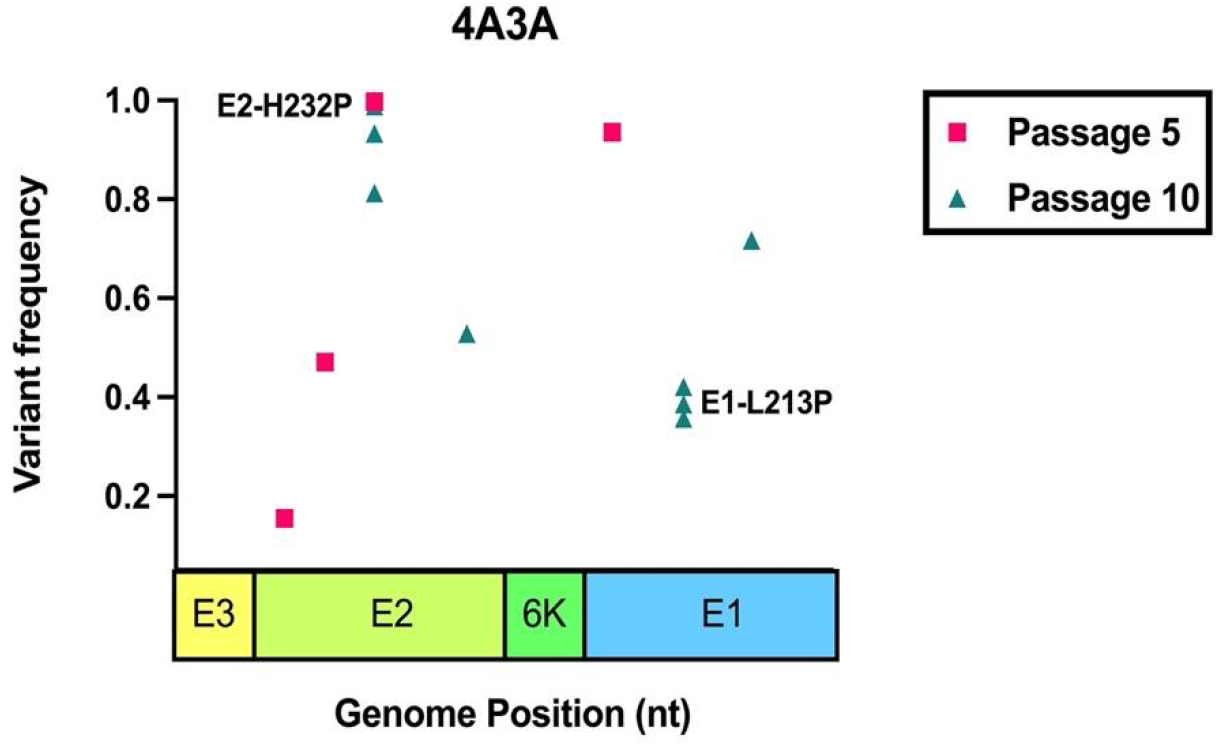
Natural evolution of Mayaro virus (MAYV) in *Anopheles gambiae* cells. Experimental evolution was performed using traditional serial passaging in 4a-3A cells. We serially passaged MAYV at a MOI of 0.01 for a total of ten passages. Following one, five, and ten passages, the viral RNA was sequenced using Illumina NGS to identify potentially adaptive mutations. No high frequency variants were identified following one passage, and as such are not depicted here.

**Figure S2:**
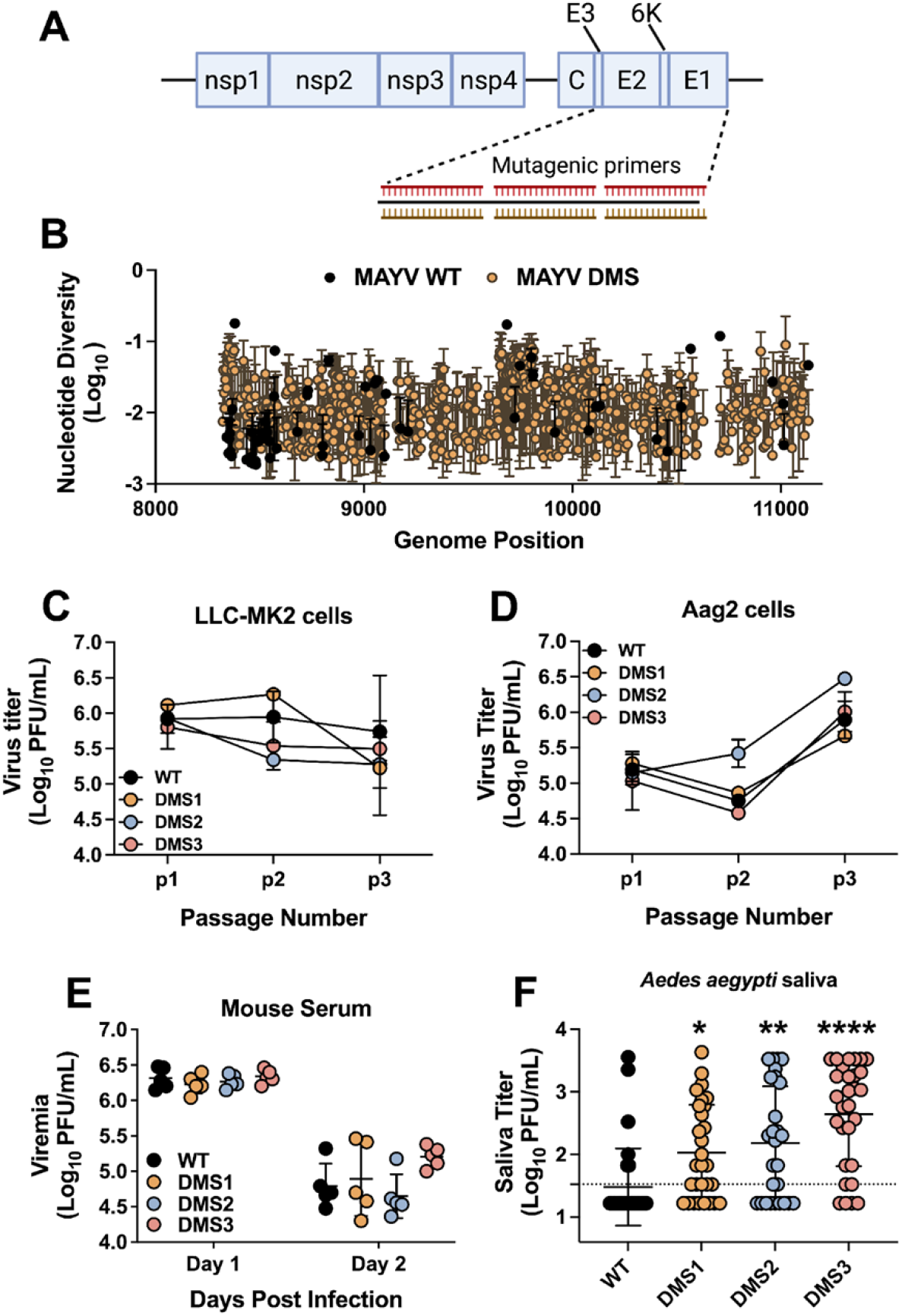
MAYV deep mutational scanning (DMS) populations are highly diverse and replicate *in vitro* and *in vivo*. **A.** Genome organization of MAYV DMS viruses. **B.** Non-synonymous nucleotide diversity for wild-type (WT) and MAYV DMS. The three independent MAYV DMS populations were combined to aid visualization. **C-D**. Three passages were performed *in vitro* at an MOI of 0.01 in LLC-MK2 (monkey kidney; **C**) and Aag2 (mosquito; **D**). **E.** Statistical analysis: * = p<0.05; ** = p<0.01; **** = p<0.0001 (one-way ANOVA with Dunnett’s correction).

**Figure S3:**
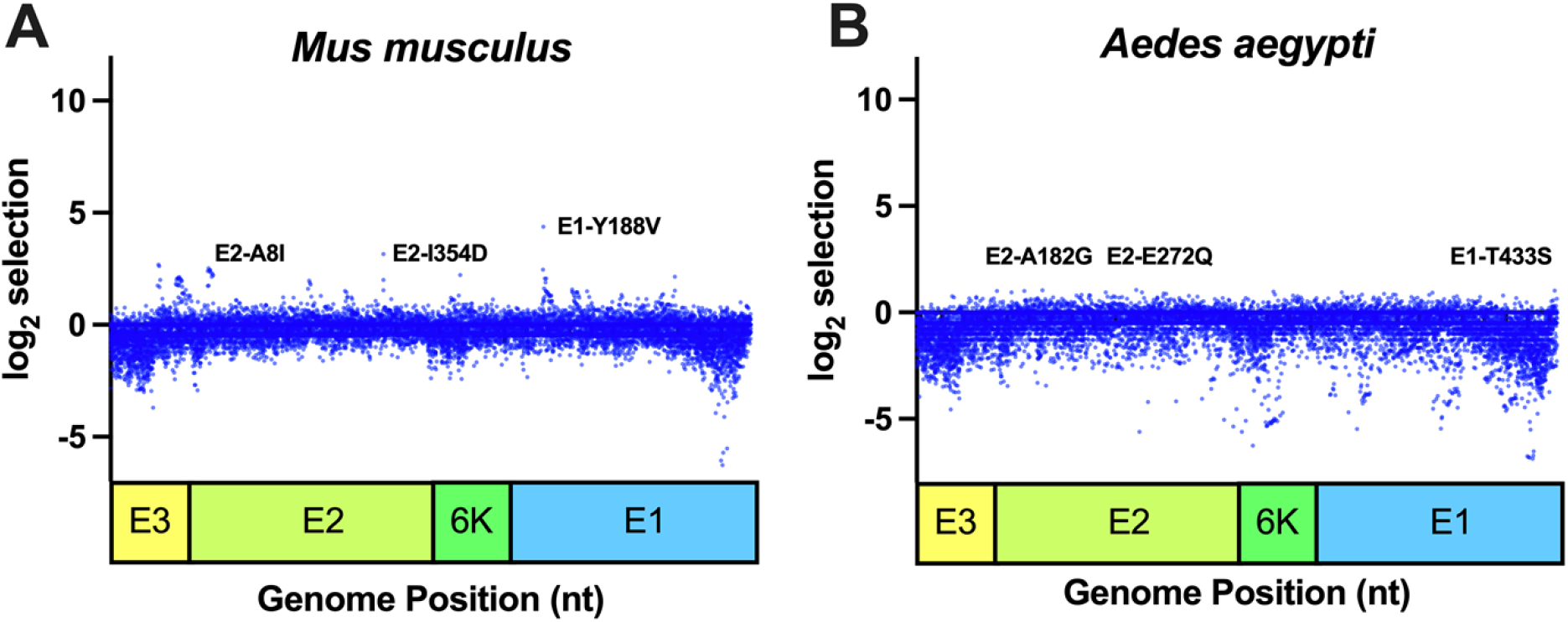
Deep mutational scanning (DMS) of Mayaro virus (MAYV) in mice and mosquitoes. **A-B.** The three MAYV DMS populations, along with WT MAYV, were used to perform three passages in mice (**A**) and *Aedes aegypti* mosquitoes (**B**). Following passage, the viral RNA was sequenced, and selection analyses were performed to identify enriched variants. The top three variants in each environment, based on selection strength, are presented for each graph.

### Adaptive mutations to *Aedes aegypti* cells come at a fitness cost

We next generated mutants identified in NGS data from both traditional passaging and DMS studies in order to measure the fitness of individual mutations (**Fig. 2A**). The fitness of mutants was measured in a high-throughput fitness screen in several cell lines against a neutral, genetically marked competitor virus, as previously described [39–41]. In Aag2 mosquito cells, two of three mutations that were enriched during Aag2 passaging (E2-T179N, p<0.0001; and E2-Q183P, p=0.0036) significantly increased viral fitness in comparison to WT (**Fig. 2B**). In U4.4, the fitness of E2-T179N was higher than WT but this difference did not reach significance (p=0.0624; **Fig. 2C**). Together, these results suggest that E2-T179N and, to a lesser extent E2-Q183P, which were identified by two distinct passaging methods, improve MAYV replication in insect cells. In LLC-MK2 cells, two of the three mutations enriched in LLC-MK2-passaged DMS populations showed significant fitness losses (E1-Y118V, p<0.0001; E2-P240F, p<0.0001; **Fig. 2D**). In fact, E1-Y118 also had lower fitness in MRC-5 human fibroblast cells, (p<0.0001; **Fig. 2E**), suggesting a mammalian-specific attenuation. We also observed fitness losses for all Aag2-enriched variants in MRC-5 (p<0.001 for all comparisons to WT), which we did not observe in LLC-MK2, suggesting a host- or cell-type specific fitness loss for these mutants. We also evaluated the fitness of mutations enriched following passage through mice and mosquitoes and observed mostly modest effects on fitness (**Fig. S4**). Notably, the E2-I354D mutant, which was enriched following passage in mice, lost fitness in both mammalian and mosquito environments.

**Figure 2.**
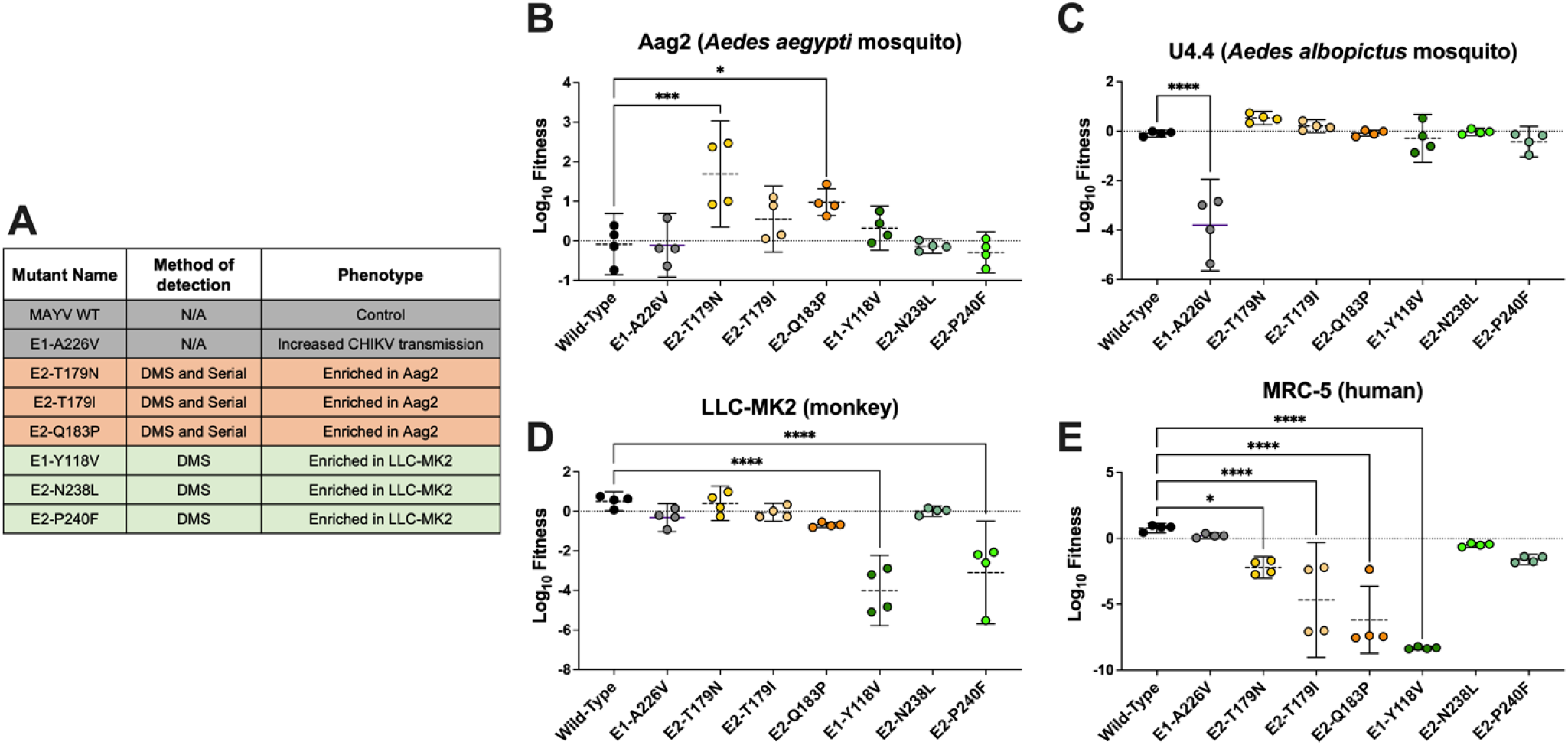
Variants enriched in Aag2 cells have increased fitness on insect cells but decreased fitness on mammalian cells. **A:** List of the viruses used for the competition assays. The method (serial passaging or DMS) used to identify the mutations, as well as the phenotypes observed for these different viruses are indicated. **B-E:** Competition assays in Aag2 (**B**), U4.4 (**C**), LLC-MK2 (**D**), and MRC-5 (**E**) cells. Cells were infected at a MOI of 0.01 using a 1:1 ratio based on PFUs for each mutant or WT with a genetically marked MAYV competitor virus. Viral supernatants were harvested at 72h post-infection for Aag2 and 48h post-infection for the other cell lines. Replication of WT and mutant viruses was assessed by RT-qPCR using specific probes labeled with different fluorophores. Log_10_ fitness was calculated by normalizing replication of each virus against a genetically marked reference virus. The mean of 4 independent experiments is represented with standard deviation. Statistical analysis: * = p<0.05; ** = p<0.01; *** = p<0.001; **** = p<0.0001 (one-way ANOVA with Dunnett’s correction).

To confirm the effect of the E2-T179N mutation on viral replication, we performed viral growth curves in Aag2 and MRC-5 cells. Results were consistent with the competition assays, with a positive effect of E2-T179N on MAYV replication in Aag2 mosquito cells (p=<0.0001 at 24h p.i.) coming at the cost of reduced replication in MRC-5 human cells (p= 0.0021 at 24h p.i.) (**Fig. 3A-B**). No differences in genome:PFU ratios were observed between WT and E2-T179N MAYV during replication in Aag2 cells (**Fig. S5**).

**Figure 3.**
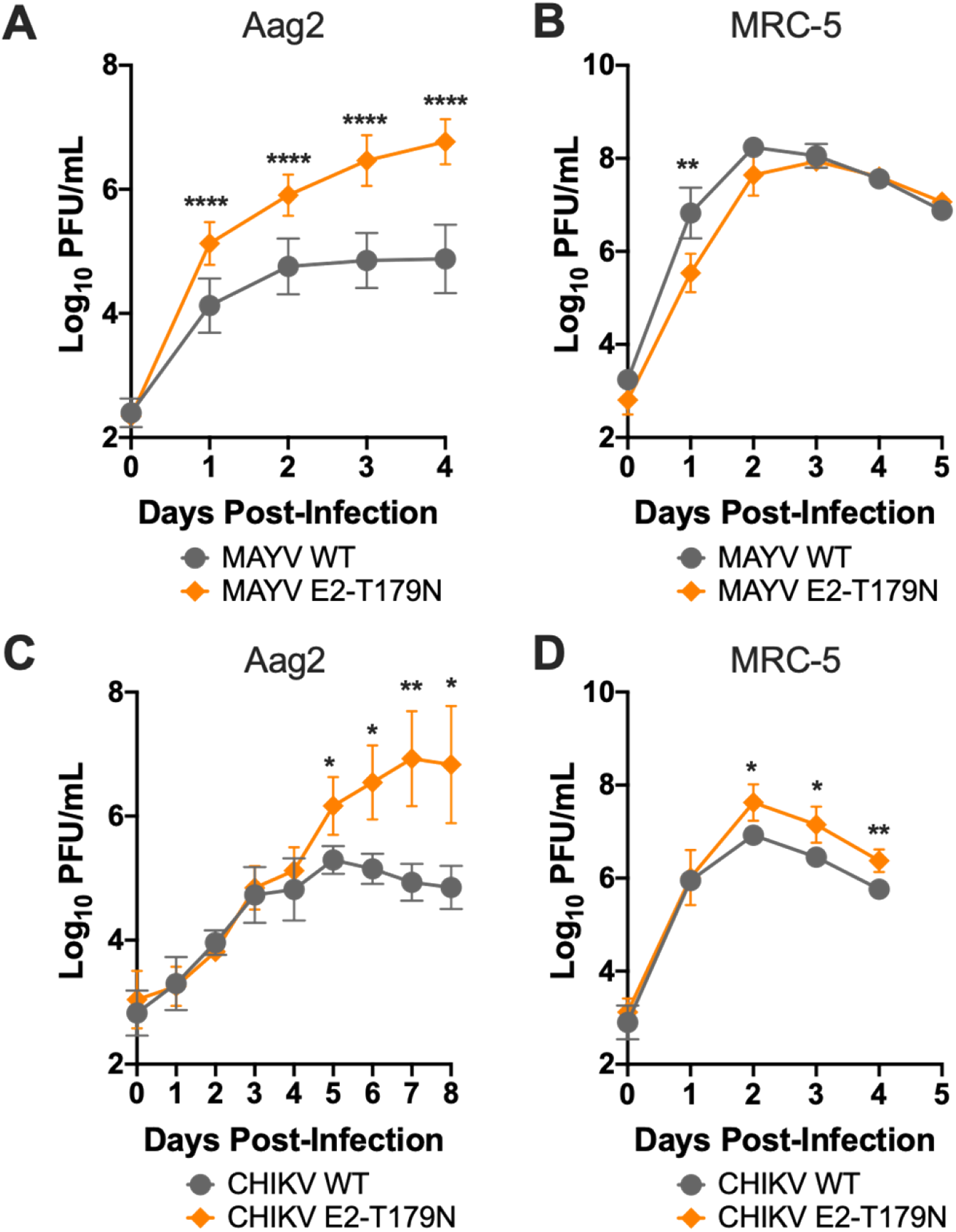
E2-T179N increases viral replication of both MAYV and CHIKV in insect cells. **A-B:** Replication of WT and E2-T179N MAYV in insect and human cells. Aag2 cells were infected with MAYV WT or MAYV E2-T179N at an MOI 0.1 (**A**), while MRC-5 were infected at an MOI of 0.01 (**B**). Replication was assessed over time by plaque assay titration. Data represents the mean of 2 independent experiments. **C-D**: Replication of WT and E2-T179N CHIKV in insect and human cells. Aag2 cells were infected with CHIKV WT or CHIKV E2-T179N at an MOI 0.1 (**C**), while MRC-5 were infected at an MOI of 0.01 (**D**). Replication was assessed over time by plaque assay titration. Data represents the mean of 2 independent experiments. Statistical analysis: * = p<0.05; ** = p<0.01; **** = p<0.0001 (two-way ANOVA using Šídák’s multiple comparisons).

Next, we assessed whether the E2-T179N mutation similarly impacts CHIKV, a closely related alphavirus that has caused major outbreaks globally in the last decades. Like MAYV, we observed that E2-T179N significantly enhanced CHIKV replication in Aag2 cells (**Fig 3C**; p=0.0108 at 120h p.i.). However, in contrast to our MAYV results, we observed an increased replication in MRC-5 cells for the mutant compared to WT CHIKV (**Fig. 3D;** p=0.0164 at 48h p.i.). Together, these results indicate that E2 position 179 is a key residue impacting viral replication not only of MAYV but also of other related alphaviruses like CHIKV, albeit with different outcomes in human cells.

**Figure S4.**
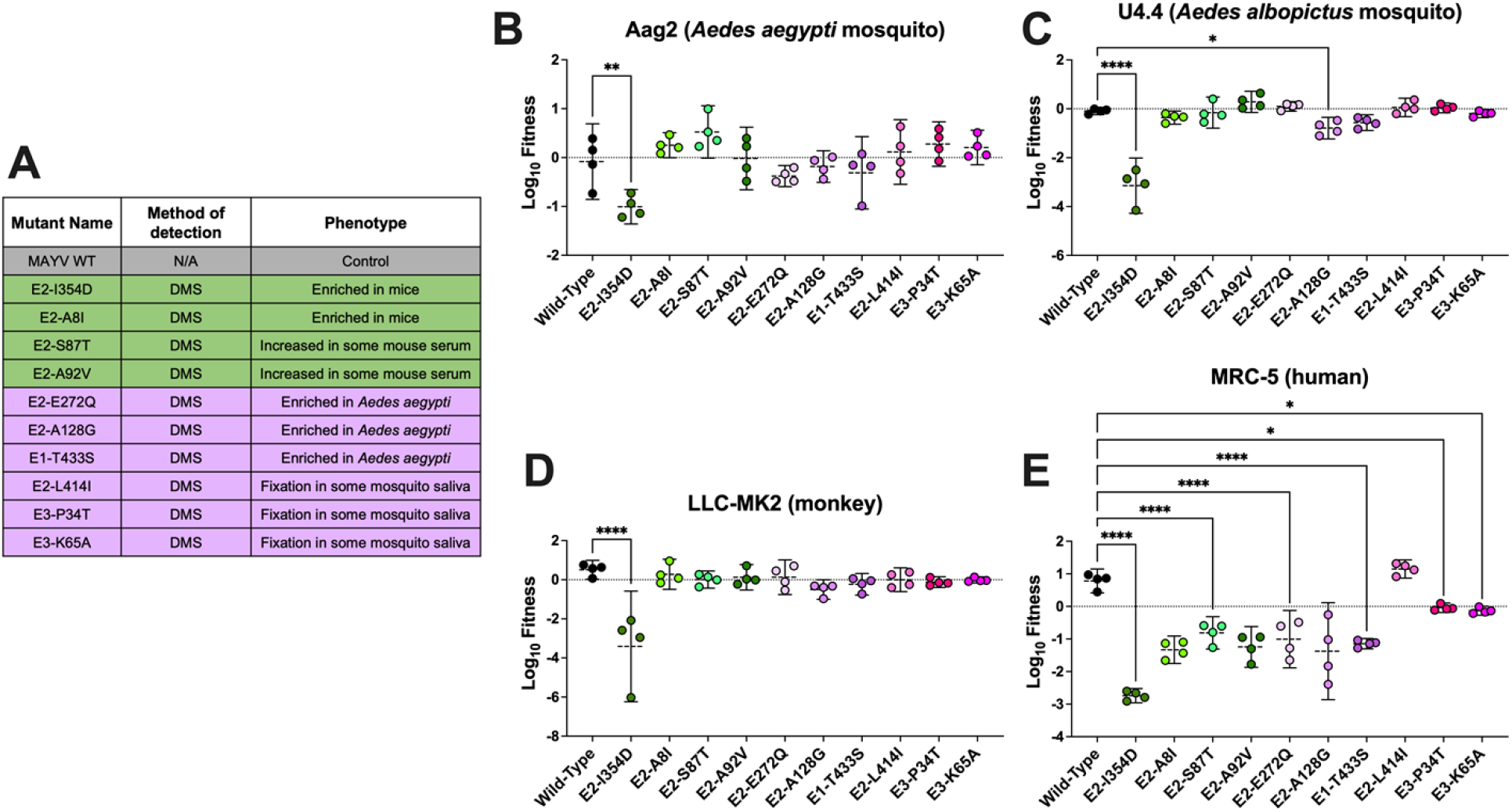
Variants enriched in mice and mosquitoes have little effect on viral fitness. **A:** List of the viruses used for the competition assays. All mutations were identified from deep mutational scanning data, and the phenotypes observed for the different viruses are indicated. The wild-type data—included as a comparison—is the same as is presented in Figure 2 in the main text. **B-E:** Competition assays in Aag2 (**B**), U4.4 (**C**), LLC-MK2 (**D**), and MRC-5 (**E**) cells. Cells were infected at a MOI of 0.01 using a 1:1 ratio based on PFUs for each mutant or WT with a genetically marked MAYV competitor virus. Viral supernatants were harvested at 72h post-infection for Aag2 and 48h post-infection for the other cell lines. Replication of WT and mutant viruses was assessed by RT-qPCR using specific probes labeled with different fluorophores. Log_10_ fitness was calculated by normalizing replication of each virus against a genetically marked reference virus. The mean of 4 independent experiments is represented with standard deviation. Statistical analysis: * = p<0.05; ** = p<0.01; **** = p<0.0001 (one-way ANOVA with Dunnett’s correction).

**Figure S5.**
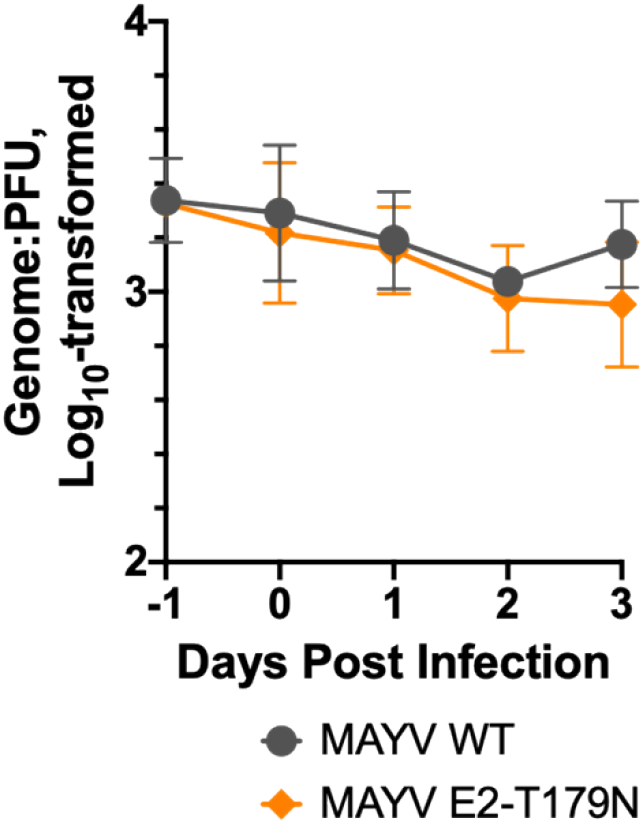
MAYV WT and E2-T179N have similar genome:PFU ratios following Aag2 infection. Aag2 cells were infected with either MAYV WT or E2-T179N at a MOI of 0.1, and genome:PFU ratios were measured each day post infection via RT-qPCR and plaque assay. The −1 day post infection represents the genome:PFU ratio of the inoculum used for the infection. Genome:PFU ratios represented are log_10_-transformed. The mean of two independent experiments is represented, error bars represent standard deviation. Statistical analysis: nonsignificant (two-way ANOVA with Šídák’s multiple comparisons test).

### E2-T179N negatively impacts cell binding independently of Mxra8

While the process by which MAYV enters insect cells remains largely unknown, there is evidence that MAYV entry in mammalian cells happens through the endosomal pathway [42] after binding with Mxra8, originally identified as the receptor for CHIKV [43,44]. Thus, we investigated whether the negative effect of E2-T179N on viral replication in human cells (**Fig. 2 and Fig. 3**) is due to decreased viral binding prior to entry and whether entry is altered via the Mxra8 receptor.

To assess viral binding, we inoculated MRC-5 cells with either WT or E2-T179N MAYV at a MOI of 0.1 and incubated the cells at 4°C to allow for binding without internalization of the virus. We then measured the amount of bound virus via RT-qPCR, which showed significantly lower relative binding of E2-T179N MAYV compared to WT MAYV (~4.3 fold less on average, p<0.0001, **Fig. 4A**). These results suggest that the attenuation of E2-T179N MAYV observed in MRC-5 cells may be a consequence of reduced binding of E2-T179N MAYV to mammalian receptors. However, when we performed the same experiment in Aag2 cells, we also observed significantly reduced binding for E2-T179N MAYV compared to WT (**Fig. S6**), although it replicated to higher levels (**Fig. 3**). This result suggests that entry pathways may be different in mammalian and insect cells, and/or that E2-T179N may act later in the viral entry process, for instance affecting E1:E2 dimer dissociation prior to endosomal fusion.

**Figure 4:**
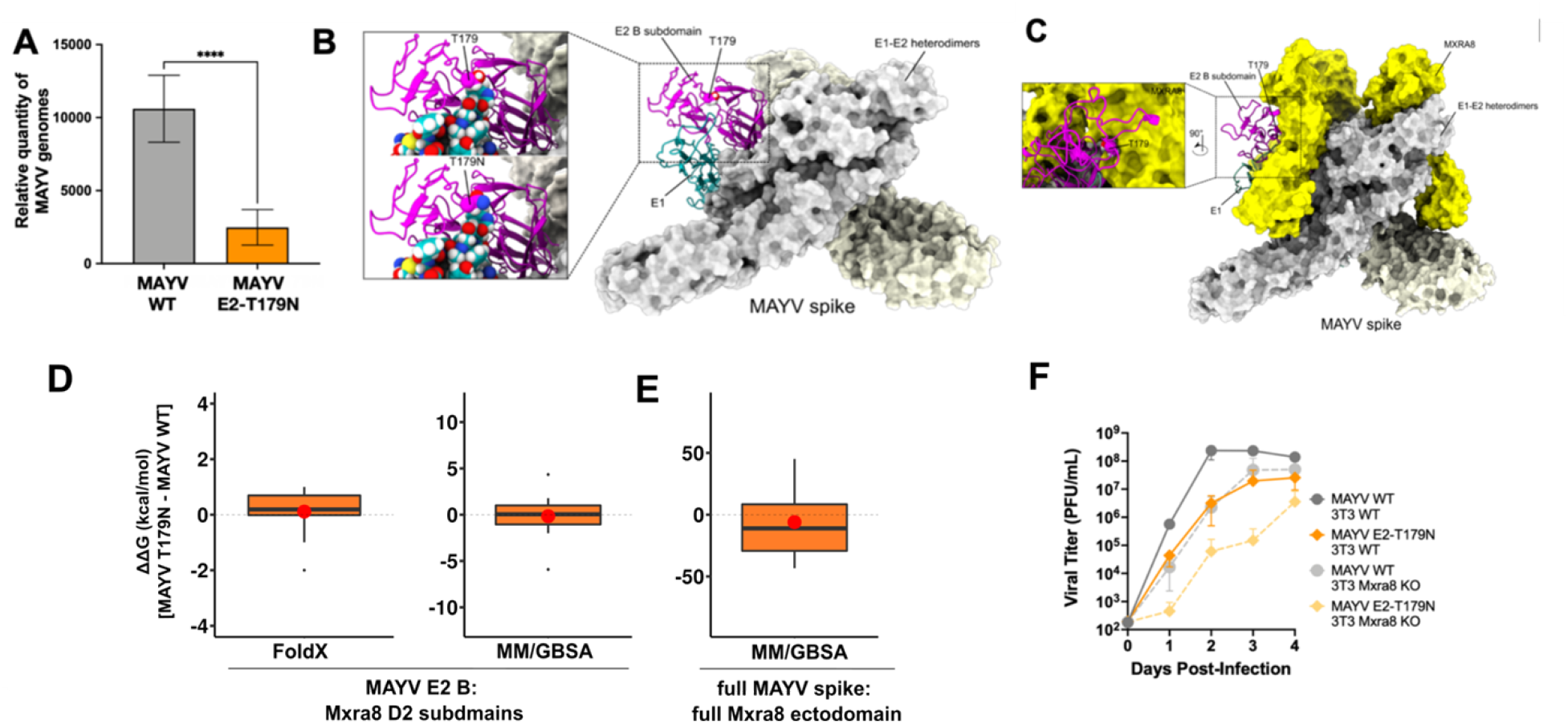
E2-T179N reduces binding to human cells independently of Mxra8. **A:** Binding of WT MAYV and E2-T179N on MRC-5 cells. Pre-chilled MRC-5 cells were infected at a MOI of 0.01, and virus was adsorbed to the cells at 4°C. Unbound virus was washed away, and bound virus was quantified by extracting RNA from the cells and performing RT-qPCR. Relative genome copies of bound virus were determined by normalizing the Ct value of bound virus to the Ct value of the housekeeping gene and virus in the inoculum. Statistical Analysis: unpaired t-test; **** = p<0.0001. Data comprise two independent binding assays with MOI of 0.1 for each virus (n=3). **B:** MAYV spike structure (PDB ID: 7KO8) composed by three E1-E2 heterodimers. Only the E1-E2 ectodomains are presented. For clarity, we showed only one heterodimer (E1 in green and E2 in magenta) in cartoon representation, the other two are represented as a surface. Zoomed-in views of T179 and the T179N mutation are shown in insets. T179, T179N and the E1 protein are represented in spheres. **C:** MAYV spike in complex with three copies of the human Mxra8 receptor (in yellow) obtained from the last frame of the 20 ns molecular dynamics simulations. For clarity, only one E1-E2 heterodimer is represented as a cartoon. Inset shows a side view of the interface between the E2 B subdomain and the Mxra8 receptor. T179 is shown in spheres. All images were generated using the ChimeraX software. **D:** Binding free energy changes (ΔΔG in kcal/mol) of 12 MAYV E2 B:Mxra8 D2 subdomain complexes (n = 12) upon MAYV T179N mutation using FoldX and MM/GBSA methods. ΔΔG was obtained by subtracting the ΔG estimated for T179N MAYV:Mxra8 interaction from the ΔG obtained for WT MAYV:Mxra8 interaction. MMGBSA calculations were performed from 2 ns molecular dynamics simulation (in triplicate) of each complex. In none of the applied methods, did the estimated ΔΔG significantly differ from zero (p > 0.05 - one sample t-test). The box plot central line indicates the median and the red circle indicates the mean. **E**: Binding free energy changes (ΔΔG in kcal/mol) of the full trimeric MAYV spike in complex with the full Mxra8 ectodomain upon MAYV T179N mutations using MM/GBSA method. The calculations were performed from a quintuplicate of 20 ns molecular dynamics simulations (n = 5) (see methods for details). ΔΔG was obtained by subtracting the relative ΔG estimated for T179N MAYV:Mxra8 interaction from the ΔG obtained for WT MAYV:Mxra8 interaction. The estimated ΔG did not significantly differ from zero (p > 0.05 - one sample t-test). The box plot central line indicates the median and the red circle indicates the mean. **F:** Growth curve of MAYV WT and MAYV E2-T179N (MOI 0.1) in 3T3 WT and Mxra8 KO lines. Experiments were performed in two independent biological experiments each performed in triplicate. A twoway ANOVA was performed on these data, and no significant differences were observed. Error bars represent the standard deviation.

To assess the potential impact of E2-T179N on MAYV structure, we used the cryo-EM structure of MAYV (PDB ID: 7KO8 [45]) aligned with the CHIKV structure in complex with Mxra8 (PDB ID: 6JO8 [46]). The E2-T179N change was not expected to have a major impact on the secondary structure of subdomain B or to impair protein-protein contacts between subdomains in the MAYV E2 (**Fig. 4B**). However, the E2-T179 position is in proximity to the binding site for the Mxra8 receptor (**Fig. 4C**). We thus reasoned that E2-T179N may impair the interaction with Mxra8, which may explain our results on viral binding (**Fig. 4A**). To investigate the E2-T179N structure and E2-T179N-Mxra8 interactions, we conducted several *in silico* analyses. First, we used two N-glycosylation prediction servers, NetNGlyc [47] and NGlycPred [48] to determine whether the E2-T179N mutation could create an additional N-glycosylation site via a N-X-S/T motif. Both methods failed to detect any additional N-glycosylation site in E2-T179N. Next, we used molecular modeling and molecular dynamics (MD) simulations with predicted free energy of binding calculations to evaluate if the E2-T179N mutation could affect the interaction between the E2 glycoprotein and the Mxra8 receptor. We obtained a ΔΔG of 0.15 ± 0.66 kcal/mol, indicating that the T179N mutation did not change predicted binding affinity to the Mxra8 D2 domain (p > 0.05, one-sample t-test) (**Fig. 4D**). To consider small adjustments in the binding of Mxra8 to MAYV ectodomains, we modeled and performed molecular dynamics simulations in quintuplicate of the full MAYV spike, formed by three pairs of E1 and E2 ectodomains, in complex with 3 copies of the entire Mxra8 ectodomain. We calculated the binding free energy using MMGBSA and computed the difference between the relative binding free energy (ΔΔG) of E2-T179N MAYV and WT MAYV spike to Mxra8. The ΔΔG predicted with MM/GBSA was −6.0 ± 26.3 kcal/mol (**Fig. 4E**), which again indicates no significant (p > 0.05, one-sample t-test) free energy change associated with the E2-T179N mutation.

To experimentally assess whether Mxra8 plays a role in the reduction of viral replication observed for E2-T179N MAYV, we infected WT and Mxra8 knock-out mouse fibroblasts (**Fig. 4F**). Our results show decreased replication of E2-T179N compared to WT MAYV, consistent with our previous results of replication attenuation in MRC-5 cells (**Fig. 3**). However, using this model, the absence of Mxra8 induced a similar reduction of viral replication of both WT and E2-T179N MAYV, confirming our *in silico* analyses showing that the reduced viral binding of E2-T179N MAYV is Mxra8-independent.

**Figure S6.**
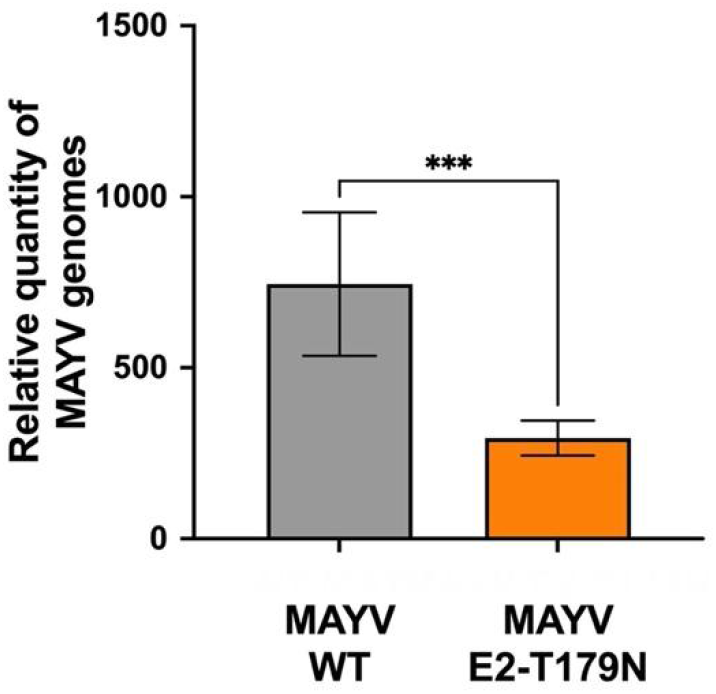
Binding affinity of WT and E2-T179N MAYV to Aag2 cells. Binding assays of MAYV WT and E2-T179N were performed by inoculating chilled Aag2 cells with virus at a MOI of 0.1 prior to adsorption at 4°C. Unbound virus was washed away, and bound virus was quantified by RT-qPCR using RNA extracted from the inoculated cells. The relative quantities of MAYV genomes were determined by normalizing to the Ct values of a housekeeping gene and the Ct value of virus in the inoculum. Data represents the mean of two independent experiments. Statistical analysis: *** = p<0.002 (unpaired t-test).

### E2-T179N increases transmissibility in the urban vector *Aedes aegypti*

Given that the E2-T179N mutation increased fitness of MAYV in Aag2 cells, we hypothesized that it may impact MAYV transmission potential in *Aedes aegypti* mosquitoes. To test this hypothesis, we exposed groups of *Aedes aegypti* to an artificial bloodmeal containing 10^6^ PFU/mL of WT or E2-T179N MAYV. To assess infection levels, we collected the midgut of infected mosquitoes, as this is the first site of viral replication. We observed that both WT and E2-T179N MAYV infected mosquitoes to high levels, with about >80% of mosquitoes readily infected (**Fig. 5A**), although the WT MAYV infection rate in mosquitoes was slightly higher than for E2-T179N (p=0.035). Similarly, we collected legs and wings to assess the ability of MAYV to disseminate to peripheral organs and did not observe significant differences between WT and E2-T179N MAYV (**Fig. 5B;** p=0.093). In contrast, we observed a significant increase in the proportion of mosquitoes able to transmit MAYV; indeed, E2-T179N MAYV could be reliably detected in the saliva of 15 to 60% of infected *Ae*des *aegypti*, while WT MAYV was detected to significantly lower levels (from 0 to 12%; **Fig. 5C**; p<0.0001). These results strongly suggest that E2-T179N dramatically increases MAYV transmission potential of *Aedes aegypti*, possibly by allowing it to infect or escape from the salivary glands and enter the saliva more efficiently.

**Figure 5.**
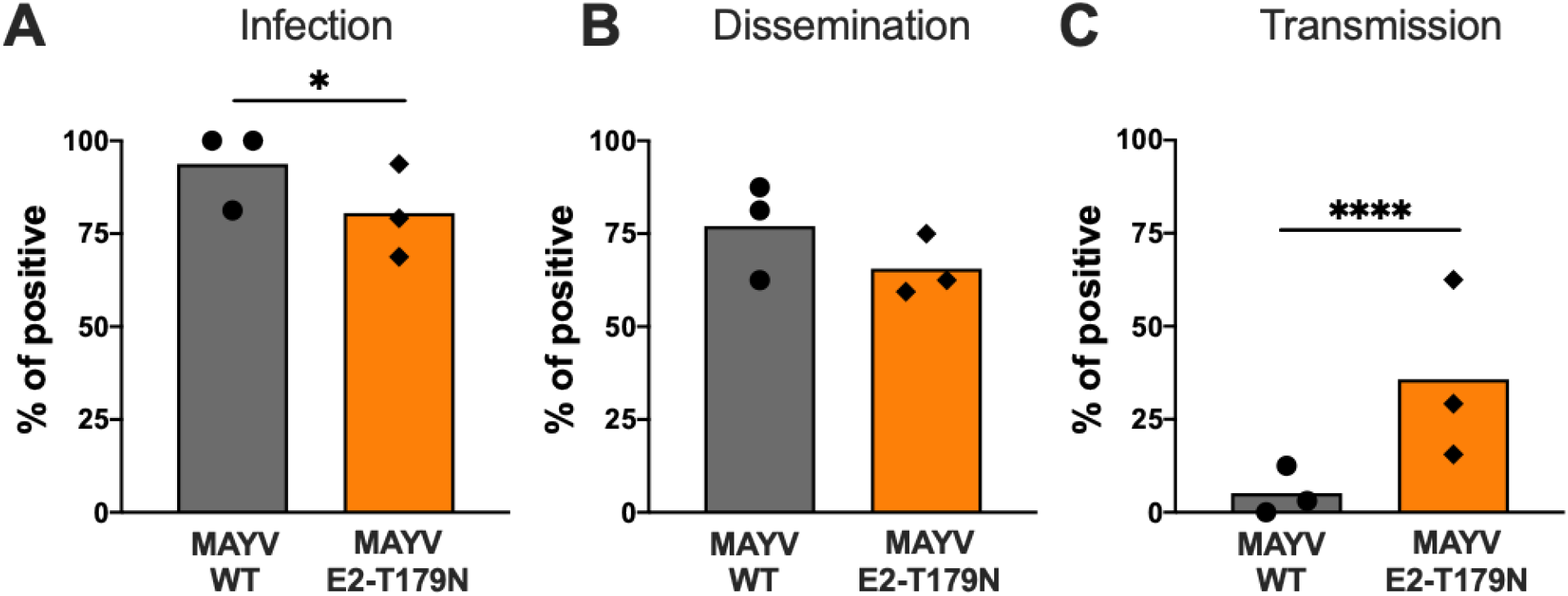
Vector competence of *Aedes aegypti* infected by WT and E2-T179N MAYV. **A:** Adult *Aedes aegypti* mosquitoes were infected with a MAYV infectious bloodmeal. Fully engorged mosquitoes were kept at 28°C with permanent access to a 10% sucrose solution. Mosquitoes were cold-anesthetized and dissected at 7 dpi. Viral titers in the midgut (**A**) and legs/wings (**B**) were quantified by plaque assay titration. Prior to dissection, saliva was harvested from live mosquitoes and viral loads were further amplified either on C6/36 or on BHK-21 cells. After 3 days, presence of virus was detected using RT-qPCR or direct visualization of CPEs (**C**). The percentage of positive midguts (**A**), legs/wings (**B**) and saliva (**C**) represent infection, dissemination and transmission rates, respectively. The mean of 3 independent experiments (with at least 16 mosquitoes each experiment; n=72 total/group) is represented. Matching symbols derive from the same experiment. Statistical analysis: * = p<0.05; **** = p<0.0001 (Two-tailed Fisher’s exact test).

### Mosquito adapted E2-T179N MAYV loses fitness in mammals

Since MAYV E2-T179N exhibited increased replication in insect cells but reduced replication in mammalian cells compared to MAYV WT, we sought to explore whether adaptation to *Aedes aegypti* comes at the cost of reduced viral fitness in a mammalian host. Thus, we infected CD-1 mice with WT MAYV and E2-T179N MAYV. Mice infected with E2-T179N MAYV had significantly lower viremia than mice infected with WT MAYV at all timepoints tested (**Fig. 6A**) but mice consistently gained weight over the course of the study in both groups (**Fig. 6B**). Mice infected with E2-T179N MAYV had significantly lower footpad swelling relative to WT MAYV at five days post infection (**Fig. 6C**; p=0.0006) and delayed peak swelling, although peak swelling between the groups was similar. To determine whether E2-T179N MAYV infection caused less tissue damage compared to WT MAYV, we collected the inoculated footpad of each mouse at either seven days post-infection and processed them for histological analysis. Based on the severity of lymphoplasmacytic myositis, scores ranging from zero to three were assigned to samples, with zero denoting normal tissue and three the greatest extent of myofiber loss across all samples. Mice infected with MAYV E2-T179N MAYV yielded significantly lower footpad scores than those infected with WT MAYV (**Fig. 6D**; p=0.0150), indicating lower levels of tissue damage at this timepoint. Images taken from the stained cross-sections of footpads show the degree of muscle fiber degeneration and immune cell infiltration, with this symptom being more severe in the group infected with WT MAYV (**Fig. 6E, panel 1**) than the E2-T179N mutant (**Fig. 6E, panel 2**). The primary infiltrating immune cells were lymphocytes, and to a lesser extent, plasma cells and macrophages. A representative image from mock-infected mice is presented in **Fig. 6E, panel 3**. These results are consistent with E2-T179N causing an attenuation of MAYV in the mammalian host

**Figure 6.**
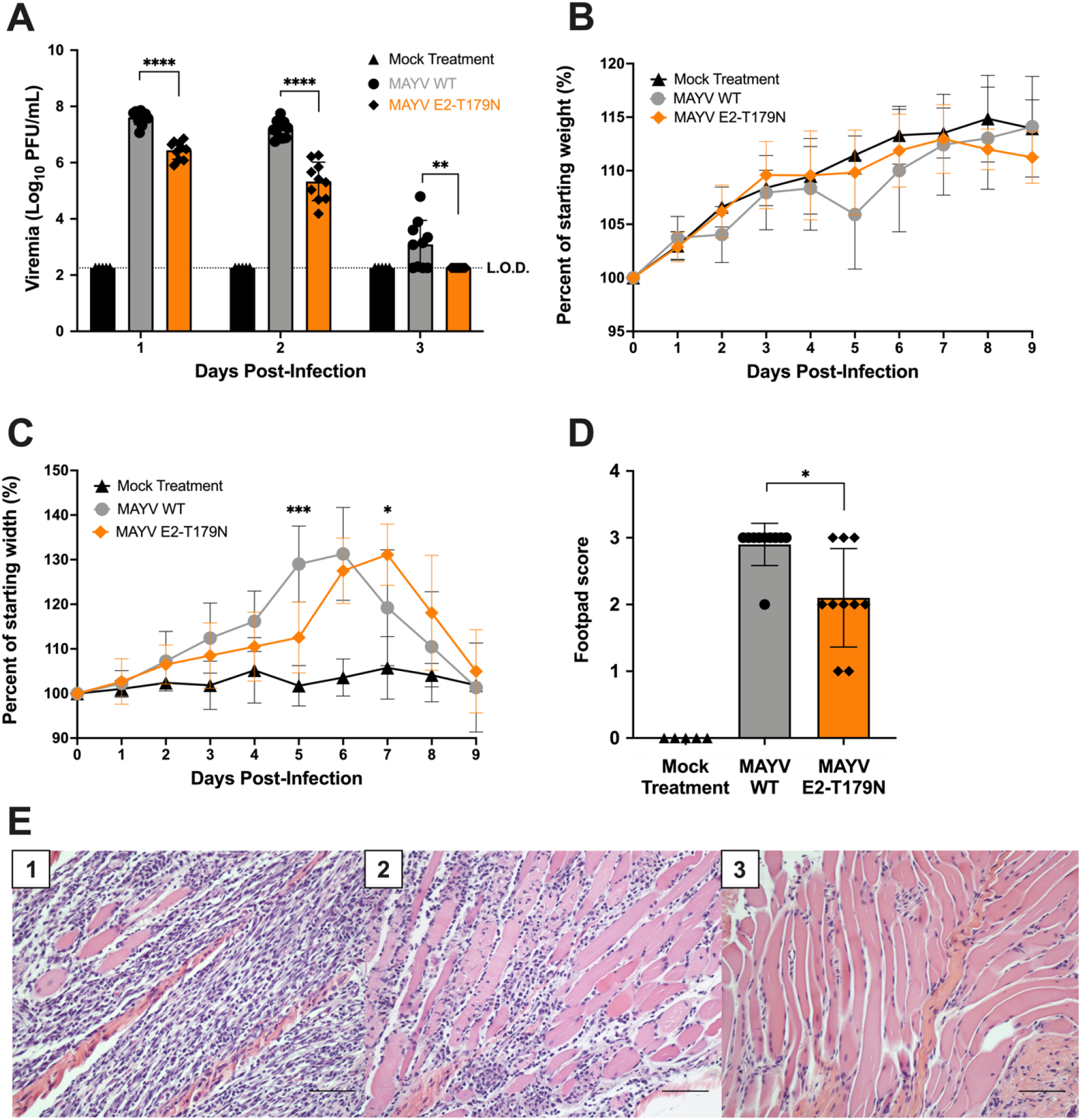
E2-T179N attenuates MAYV replication and disease in mice. Four-week-old CD-1 mice were infected with 10^5^ PFU in 50 μL via injection to the left, hind footpad. **A:** Viral load in serum was assessed daily by plaque assay. Statistical comparisons were made using multiple unpaired t tests with correction for multiple comparisons using the false discovery rate method of Benjamini, Krieger, and Yekutieli. **B**: Mice were weighed daily following infection to assess weight change. **C:** Footpad swelling was measured daily using a digital caliper. **B-C:** Statistical comparisons were made using a two-way ANOVA with Dunnett’s correction for multiple comparisons. Data represent the percent of the weight or footpad width before infection. **D:** Footpad swelling score as determined by histopathology whereby a score of 0 indicates a normal state, 1 mild (<25%) myofiber loss, 2 moderate (25-50%) myofiber loss, and 3 severe (>50%) myofiber loss. Statistical comparisons were made by Mann-Whitney test. **E:** Hematoxylin and eosin stain of muscular tissue in left, hind footpad of mice seven days post infection with WT (panel 1) or E2-T179N (panel 2) MAYV or after a mock treatment (panel 3). The scale bar in the images is 200 microns. Studies were performed in two independent experiments, using 10 mice per infected group and five mock-treated mice. Error bars represent standard deviation from the mean.

## Discussion

MAYV is an emerging viral threat with epidemic potential. The main goal of this study was to determine whether evolution in MAYV may lead to viral emergence, particularly through adaptation to the urban vector *Aedes aegypti*. We identified E2-T179N as a mutation that confers increased replication in *Aedes aegypti* cells (**Fig. 2-3**) and enhances viral transmission potential by *Aedes aegypti* mosquitoes (**Fig. 5**). Thus, we confirm that MAYV has the potential to better adapt to *Aedes aegypti* and therefore continues to represent a risk to human health.

Fortunately, the same E2-T179N mutation caused a fitness reduction in human cells (**Fig. 2-3**), suggesting that this mutation alone would not favor urban MAYV transmission. Other roadblocks remain in the way of MAYV emergence: notably, it is still unclear whether the viremia induced by MAYV in humans is sufficient to sustain human-to-human amplification. Serum viral loads of 10^4^-10^5^ PFU/mL MAYV have been observed in a study involving 21 humans with confirmed MAYV infection during the three days following symptom onset [4]. In comparison, infectious titers for CHIKV range from 10^3.9^ to 10^6.8^ PFU/mL [49]. Thus, titers in humans appear to be slightly lower for MAYV than the successful human pathogen, CHIKV, which may limit emergence potential without further adaptation. Furthermore, the geographical overlap between MAYV and CHIKV may prevent MAYV emergence, as studies in mice and have suggested that a prior CHIKV infection confers cross-protection against MAYV infection [52,53].

The two different experimental evolution approaches we used (serial passaging and DMS) lead to the identification of E2-T179N as a key mutation involved in MAYV replication in *Aedes aegypti* cells. Notably, we observed that only three serial passages of DMS viruses in Aag2 allowed identification of this adaptive mutation (**Fig. 1F**), while we observed this mutation in only 50% of the replicates in our serial passaging experiment after ten passages (**Fig. 1D**). This confirms that DMS passaging speeds up the process of selection, although DMS can only be targeted to a portion of the viral genome. Thus, we conclude that combining these two approaches represents an efficient strategy to identify adaptive mutations with high confidence. Perhaps unsurprisingly, most mutations detected with both approaches occurred within the receptor-binding (E2) protein, highlighting the importance of this protein in dictating cell tropism. Furthermore, the residues with the strongest impact on fitness (positions 179 and 183 in E2) both reside within domain B, which we previously showed was critical for cell tropism [51]. A surprising finding in our DMS studies was that variants enriched during passage in LLC-MK2 either had neutral or deleterious impacts on fitness in mammalian cells. However, these data are consistent with results observed in ZIKV [38] where mammalian fitness gains were only observed with a double mutant. A possible explanation for this is the importance of cooperative interactions with other residues (either on the same genome or on other genomes) occurring within the viral population. Moreover, a mutation that was slightly enriched during mouse passage (E2-I354D) decreased fitness appreciably in both mammalian and insect cells; thus, this mutation represents a broadly attenuating mutation and suggests that DMS studies may be useful for developing rationally designed attenuated vaccine viruses. For example, in future studies, selecting the residues most differentially selected against may lead to a highly attenuated virus and could help increase understanding of fundamental virus biology using a safer loss-of-function approach.

The mechanisms underlying the impact of E2-T179N on MAYV replication in *Aedes aegypti* and human cells are still unclear. We hypothesize that this represents an evolutionary trade-off: because of its dual-host nature, MAYV cannot optimize its replication in both insect vectors and mammalian hosts simply through the E2-T179N mutation, possibly due to the need to use different receptors in each host. However, it is possible that additional mutations in E1 or E2 may compensate for this defect, and/or that evolution at the T179 towards other residues may confer increased replication in both hosts. Interestingly, we observed that two distinct mutations (T179N and T179I) were enriched in mosquito cells. Therefore, testing other T179 mutation combinations may provide additional insights into MAYV viral evolution potential. Interestingly, E2-T179N CHIKV similarly displayed increased fitness in *Aedes aegypti* cells (**Fig. 3C**), but, in contrast to MAYV, also showed increased fitness in human fibroblasts (**Fig. 3D**). Thus, this residue is important for fitness for both viruses but the differential impact on fitness suggests different mechanisms or epistatic interactions. This result is consistent with the effect we observed with the E1-A226V MAYV mutant, which is known to enhance CHIKV’s fitness in *Aedes albopictus* [54] but greatly reduced MAYV’s fitness in U4.4 cells (**Fig. 2C**). Given that E2-T179N in MAYV did not alter fitness in monkey or *Aedes albopictus* cells (**Fig. 2B-C**), we cannot conclude that this is a mammalian- or insect-specific phenotype; rather, it appears to be cell type specific. Similarly, it is unlikely that temperature is mediating the phenotypes we observed because we would expect similar results in LLC-MK2 and U4.4 as we observed in MRC-5 and Aag2. It is possible that this mutation alters receptor binding in a cell-specific manner; although we did not observe differences in growth rates in cells lacking the Mxra8 receptor or differences in predicted free energy of MAYV binding to Mxra8 upon T179N mutation (**Fig. 4**). Given that CHIKV and closely related viruses still replicate to high titers in the absence of Mxra8 [43], it is possible that this mutation disrupts interactions with other, yet to be discovered, cell surface receptors that may be different in insect and human cells. Other possible explanations for the phenotypes observed are disrupted E1:E2 interactions, since this residue resides in domain B, which interacts with E1. Future studies will help shed light on the mechanisms responsible for the phenotypes observed.

In addition, the E2-T179N mutation led to attenuation *in vivo* in a mouse model (**Fig. 6**). Future work will establish whether attenuation of MAYV-induced disease is solely due to lower viral replication, or to other immune parameters such as inflammation, cytokine response, IFN signaling, etc. Furthermore, future studies will assess whether viremia levels in mice are still sufficient to cause productive infection in mosquitoes. Our work suggests that mutations in the E2 of MAYV may favor its dissemination through the usage of alternate mosquito vectors and should therefore be monitored closely with appropriate surveillance programs.

## Material and Methods

### Cells, viruses and plasmids

Vero, BHK-21, LLC-MK2, and MRC-5 cells were grown in Dulbecco’s modified Eagle’s medium (DMEM, Gibco), containing 10% fetal calf serum (FBS; Gibco), 1% penicillin/streptomycin (P/S; Thermo Fisher) in a humidified atmosphere at 37°C with 5% CO_2_. U4.4 and Aag2 cells were maintained in Leibovitz’s L-15 medium (Gibco) with 10% FBS, 1% P/S, 1% non-essential amino acids (Sigma) and 1% tryptose phosphate (Sigma) in a dry atmosphere without CO_2_ at 28°C. 3T3 WT and Mxra8 KO cells were a kind gift of Dr. Mike Diamond and were maintained at 37°C with 5% CO_2_ in DMEM (Genesee Scientific) supplemented with 10% fetal bovine serum, 1% non-essential amino acids, 50 μg/mL gentamicin sulfate, and 25 mM HEPES. For some studies, Aag2 were maintained in Schneider’s insect medium (Genesee Scientific) with 7% FBS, 1% non-essential amino acids, 2.5 μg/μL Amphotericin B, 50 μg/mL gentamicin sulfate, and 5% tryptose phosphate broth without CO_2_ at 28°C. MAYV strain TRVL 4675 was derived from an infectious clone we previously described that contains an SP6 promoter to generate infectious RNA [55]. To generate the MAYV infectious clones under the CMV promoter, we amplified the entire MAYV genome from the previously described SP6-containing plasmid along with the pcDNA3.1-based CMV vector [41] containing overlapping ends. All clone sequences were confirmed using next generation sequencing (NGS). The generation of the infectious molecular clone for CHIKV strain SL-CK1 was described previously [41].

### Viral stocks

To generate MAYV viral stocks for the serial passaging experiments, a T25 flask of BHK-21 cells at 75% confluency was transfected with 10 μg of MAYV CMV-promoter driven plasmid using TransIT-LT1 Transfection Reagent (Mirus) according to the manufacturer’s instructions. Viral supernatants were collected 48h later and used to perform one blind passage on C6/36 cells. The high titer viral stock used for viral growth curves and mosquito experiments was generated the same way, except that the virus was passaged twice on Vero cells (MOI 0.01). The generation of MAYV and CHIKV mutant stocks was performed by transfecting 500 ng of DNA into a 24-well plate well using JetOptimus (Polyplus, France). The mutants were then passaged once in BHK-21 cells at an MOI of 0.01. Viral stocks were titrated by plaque assay.

### Plaque assay

Vero cells were seeded in 24-well plates (10^5^ cells per well) and infected with serial dilutions of infectious supernatant diluted in RPMI-1640 with 2% FBS and 10 mM HEPES. After 1h at 37°C, a semi-solid overlay consisting of 1.5% methylcellulose, 2X EMEM, 4X L-glutamine, 0.735% sodium bicarbonate, 0.2 mg/mL gentamicin sulfate, 4% heat-inactivated FBS, and 20 mM HEPES was added to the cells. Cells were fixed with 4% formalin solution, pH 6.2 at 3 or 4 dpi and stained with 0.1% crystal violet.

### Viral growth curves

Two days prior to infection, cells were seeded in a 24-well plate and infected at 60-80% confluency. Cells were infected in triplicate at a MOI of 0.01 for all cells except Aag2, which were infected at MOI 0.1. Virus was diluted in RPMI-1640 (Genesee Scientific) media containing 2% FBS and 10 mM HEPES. One hour post-infection, viral inoculum was removed, cells were washed once with PBS, and the appropriate culture media was added to the cells. Cells were incubated at either 37°C or 28°C, according to cell type and conditions listed above. Supernatant was collected every 24 hours for the indicated time points and frozen until viral titration.

### Serial passaging

We infected cells at a MOI of 0.01 as indicated above for viral growth curves, except infections were carried out with 6 replicates instead of 3. Viral supernatants were collected (2 days p.i. for BHK-21 cells; 3 days p.i. for U4.4 and Aag2 cells) aliquoted, and frozen. After each passage, all supernatants were titrated by plaque assay. Supernatants collected at passages 1, 5 and 10 were sequenced using NGS as described below.

### NGS for serial passaged samples

RNA of 100 μL of each sample supernatant was extracted using TRIzol reagent (Invitrogen) following the manufacturer’s protocol. RNA was eluted in 30 μL of nuclease-free water. After quantification using Quant-IT RNA assay kit (Thermo Fisher Scientific), viral RNA was enriched using polyA selection NEBNext Poly(A) mRNA Magnetic Isolation Module (NEB). Libraries were prepared with NEBNext Ultra II RNA Library Prep Kit for Illumina. The quality of the libraries was verified using a High Sensitivity DNA Chip (Agilent) and quantified using the Quant-IT DNA assay kit (Thermo Fisher Scientific). Sequencing of the libraries was performed on a NextSeq 500 sequencer (Illumina) with a NextSeq Mid Output kit v2 (151 cycles).

### Generation of MAYV envelope DMS populations

DMS populations were created in the genes encoding E3, E2, 6K, and E1 using the SP6-driven MAYV infectious clone as previously described [36-38] with modifications. Notably, we used a bacteria-free cloning approach that we have previously described [56,57]. DMS mutagenesis primers are listed in **Supplementary File 4**. The forward mutagenesis primers pool was used with a corresponding MAYV reverse end primer (5’ CCCGCATTACACGGTACTTATGAT 3’) The reverse mutagenesis primers pool was used with a corresponding MAYV forward end primer (5’ CGGAAGGCACAGAGGAGTGG 3’). We performed one round of mutagenesis with ten PCR cycles. All PCRs were performed with SuperFi II PCR master mix (Invitrogen). The fragments were then joined by PCR. The vector containing the remaining MAYV genome was amplified to create overlapping ends with the mutagenized fragments that are compatible with Gibson assembly. The primer sequences used to amplify the vector are forward (5’ CCACTCCTCTGTGCCTTCCG 3’) and reverse (5’ ATCATAAGTACCGTGTAATGCGGG 3’). Amplicons were run on a 0.6% agarose gel containing GelGreen nucleic acid stain, excised, and then purified using the NucleoSpin Gel and PCR cleanup kit (Macherey-Nagel). Amplicons were then assembled (1:1 insert:vector molar ratio) into circular molecules using the NEBuilder HiFi DNA Assembly Master Mix incubated at 50°C for two hours. To confirm that no parental plasmid vector was carried through the process, for each mutant, we included a control containing the DNA fragments but no assembly mix; this was then treated identically to the other samples for the remainder of the process. The assembly was then digested with exonuclease I, lambda exonuclease, and DpnI (all from NEB) to remove singlestranded DNA, double-stranded DNA, and bacterial-derived plasmid DNA, respectively. This product was then amplified by rolling circle amplification (RCA) using the Repli-g mini kit (Qiagen). The RCA product was linearized with SgrAI (NEB) and purified. Capped RNA was generated using the mMESSAGE mMACHINE SP6 kit (Invitrogen) and then transfected into a T150 flask of BHK-21 cells using JetMessenger (Polyplus). Viral supernatants were harvested 48h later, clarified by centrifugation, and then precipitated using polyethylene glycol (PEG) 8000 [58]. We constructed three replicate libraries by performing all the steps independently for each replicate starting with individual PCR reactions.

### DMS passaging

Passaging *in vitro* was performed in Aag2 and LLC-MK2 at a MOI of 0.01 for a total of three passages. Following one passage, the virus was titrated by plaque assay and the next passage was initiated at the same MOI. Only one passage was performed for *in vivo* studies. Three week-old female C57BL/6J mice (Jackson laboratory) were inoculated with 10^6^ PFU of MAYV WT or DMS populations in the left hind footpad. The mice were bled on days one and two post-infection to measure viremia and collect virus for sequencing. *Aedes aegypti* females originally collected in Guerrero (Mexico) were exposed to an infectious bloodmeal containing 3×10^7^ PFU/mL. Fully bloodfed mosquitoes were separated and allowed to incubate at 28°C for ten days. Following incubation, salivary secretions were collected in viral diluent containing RPMI-1640 media supplemented with 25 mM HEPES, 1% BSA, 50 μg/mL gentamicin, and 2.5 μg/mL amphotericin B. We collected only saliva samples because this best represents the virus with potential to transmit to the mammalian host.

### DMS sequencing and analysis

Viral RNA for sequencing was extracted using the Direct-zol RNA extraction kit (Zymo Research). RNA was converted to cDNA using the Maxima H Minus cDNA Synthesis Master Mix (Invitrogen). PCR amplicons containing the DMS region were amplified using SuperFi II master mix with MAYV specific primers: forward (5’ AGTGGGTAAGCCTGGCGACA 3’) and reverse (5’ TAAATCGGTCCGCATCATGCAC 3’). The fragments were purified using a 1x ratio of AMPure XP (Beckman). Libraries were prepared using Nextera XT and sequenced on an Illumina NextSeq 500. Data was analyzed using the dms_tools2 software, which is available at https://jbloomlab.github.io/dms_tools2/. Differential selection was calculated by comparing the starting virus stock to the post-passage samples.

### NGS data analysis

Raw sequencing data was deposited under the Sequence Read Archive (SRA, NCBI) under bioproject number PRJNA796940. The raw NGS reads were first trimmed using BBDuk, a tool from the BBMap tool kit, to remove the sequencing adapters and reads with a quality score of less than 30 [59]. The sequences were then aligned to the MAYV genome using the Burrow Wheeler Aligner (BWA) tool with no added parameters [60]. The resulting .bam files were then sorted using Sambamba [61]. Indels and variants were called using LoFreq [62] and then filtered to remove variants present at less than 5% frequency. We then used snpdat to annotate the variants [63]. A variant threshold was calculated by comparing the coverage depth to the allele frequency. Specifically, we took the reciprocal frequency of a given variant and multiplied by 10; if the sequencing coverage at this site was greater than that number, it was considered positive; otherwise, it was discarded [64].

### Site-directed mutagenesis and virus rescue

All mutants were created using our bacteria-free cloning approach described above with some modifications. The CMV-promoter-driven plasmids for MAYV and CHIKV were used to make mutants. Mutations were incorporated into PCR primers that created 20-30 bp overlaps for Gibson assembly; all primer sequences are available upon request. PCR fragments were amplified using Platinum SuperFi II PCR master mix (Invitrogen). The assembled product was amplified by RCA with the FemtoPhi DNA amplification kit (Evomic Science). Virus rescue was performed by transfecting RCA products directly into BHK-21 cells using JetOptimus (Polyplus), as we have previously described [65]. Viral titers were measured by plaque assay. To sequence the viral genome, we extracted RNA, performed DNase treatment to remove residual plasmid DNA, and generated cDNA using Maxima RT (ThermoFisher). PCR amplicons were generated using SuperFi II PCR master mix. Following gel or PCR purification, amplicons were submitted for Sanger sequencing at the Virginia Tech Genomics Sequencing Center.

### Genome:PFU ratio

A viral growth curve was performed in Aag2 cells, as above, with a MOI of 0.1 for both MAYV WT and E2-T179N. Supernatant was harvested each day post infection and titrated by plaque assay. Viral genomes were quantified each day post infection via RT-qPCR on a Bio-Rad CFX96 Touch Real-Time PCR Detection System. Briefly, supernatant was diluted 1:5 in nuclease-free water before performing reverse transcription using the New England Biolabs Universal One-Step RT-qPCR kit followed by SYBR green-based qPCR using the following primers: MAYV 5028For. (5’ CCTCTGTTAGTCCTGTGCAATAC 3’); MAYV 5108Rev. (5’ AAGGTGCTTAGGGAGCTACT 3’). A standard curve was generated using full-length MAYV genomic RNA derived from a SP6-containing MAYV infectious clone [65] to calculate MAYV genomes per milliliter of supernatant, and the genome:PFU ratio was calculated by dividing the genome concentration by the PFUs per milliliter of supernatant achieved via plaque assay.

### Competition assays

*In vitro* competition assays were performed against a genetically marked reference virus essentially as previously described [39-41]. Briefly, the competitor and test virus (in this case WT or mutant MAYV) were mixed at an equal PFU:PFU ratio and then used to infect cells at a MOI of 0.01. Virus-containing supernatant was harvested at two days p.i. for all cell lines used: Aag2, U4.4, MRC-5 and LLC-MK2. The proportion of each virus was measured using TaqMan RT-qPCR with probes specific to each virus containing ZEN / Iowa Black FQ quenchers. The primers used are as follows: MAYV 5028For. (5’ CCTCTGTTAGTCCTGTGCAATAC 3’); MAYV 5108Rev. (5’ AAGGTGCTTAGGGAGCTACT 3’); WT probe (5’ FAM CACAGTGAAACTACTGTAAGCTTGAGCTCG 3’); Competitor probe (5’ JOE CACAGTGAAACTACTGTttcCcTttcCTCG 3’). The genome copies of each virus in a given sample were calculated using a standard curve of viral RNA derived from the SP6 promoter-based clones for both WT and the marked reference virus. The relative fitness was determined with methods previously described [39,66] using the genome copies for each virus. Briefly, the formula *W=[R(t)/R(0)]^1/t^* represents the fitness (W) of the mutant genotype relative to the common competitor virus, where *R(0)* and *R(t)* represent the ratio of mutant to competitor virus in the inoculation mixture and at *t* days post-inoculation, respectively.

### Binding assays

Pre-chilled MRC-5 or Aag2 cells were washed with 4°C PBS containing 5% w/v BSA prior to inoculation with WT MAYV or MAYV E2-T179N at a MOI of 0.1. Adsorption of the virus proceeded for 30 minutes at 4°C. Inoculum was removed, and cells were washed six times with the 4°C PBS solution before RNA extraction. New England Biolabs Universal One-Step RT-qPCR kit was used to prepare cDNA from total RNA. MAYV genomes in the total RNA were quantified through SYBR green-based RT-qPCR on a Bio-Rad CFX96 Touch Real-Time PCR Detection System using the following primers: MAYV 5028For. (5’CCTCTGTTAGTCCTGTGCAATAC 3’); MAYV 5108Rev. (5’ AAGGTGCTTAGGGAGCTACT 3’); GAPDHFor. (5’ CCAGGTGGTCTCCTCTGACTT 3’); GAPDHRev. (5’ GTTGCTGTAGCCAAATTCGTTGT 3’); rp49For. (5’ AAGAAGCGGACGAAGAAGT 3’); rp49Rev. (5’ CCGTAACCGATGTTTGGC 3’). The housekeeping primers for GAPDH and rp49 were used with MRC-5-derived RNA and Aag2-derived RNA, respectively. Relative MAYV genomes were calculated by normalizing the Ct values of MAYV to the inoculum and the housekeeping gene. An unpaired t-test was performed on these data spanning two biological replicates each for MRC-5 and Aag2 cells.

### Mouse infection

Four-week-old, female CD-1 mice were inoculated with 10^5^ PFU/mL of either WT MAYV or MAYV E2-T179N in 50 μL of RPMI-1640 via injection to the left, hind footpad; in the same manner, a mock-infected group was injected with RPMI-1640. At days 1-3 p.i. serum was collected via submandibular bleed and titrated for viremia. Weights of the mice were taken and footpad widths were measured using digital calipers daily following infection. At seven d.p.i. a subset of mice was euthanized for collection of the left, hind footpad for histopathology. Footpad tissue was fixed in 4% formalin solution, pH 6.2 before decalcification in 10% EDTA pH, 7.3 at 4°C. Tissues were hematoxylin- and eosin-stained, imaged and assigned scores by a board-certified pathologist according to the extent of myofiber loss. The remaining mice were euthanized at 9 d.p.i., and footpads were collected and processed as mentioned above.

### Mosquito rearing, feeding and titration

Laboratory colonies of *Ae*des *aegypti* were established from field collections in Kamphaeng Phet Province, Thailand. All the experiments were performed within 20 generations of laboratory colonization. The insectary conditions for mosquito maintenance were 28°C, 70% relative humidity, and a 12h light: 12h dark cycle. Adults were maintained with permanent access to a 10% sucrose solution. Adult females were offered commercial rabbit blood (BCL) twice a week through a membrane feeding system (Hemotek). 6-8 days old female mosquitoes were selected and starved overnight prior to the bloodmeal. They were then fed with an infectious bloodmeal consisting of 2 mL of previously washed human blood (ICareB platform, Institut Pasteur), 1 mL of MAYV viral solution (3×10^6^ PFU diluted in L15 media with P/S, NEAA, 10% FBS and 1% sodium bicarbonate) and 5 mM ATP (Sigma). Mosquitoes were offered the infectious bloodmeal for 20 min through a membrane feeding system (Hemotek) set at 37°C with a piece of pig intestine as the membrane. Following the bloodmeal, fully engorged females were counted, selected, and kept at 28°C in 70% relative humidity and under a 12 h light: 12 h dark cycle with permanent access to 10% sucrose. 7 days after the bloodmeal, mosquitoes were cold-anesthetized and dissected. Legs and wings were collected on ice. Mosquitoes were salivated for 20 min in a tip containing 20 μL of FBS (Gibco), and midguts were collected. Body parts were collected in microtubes (Qiagen) containing 5 mm diameter stainless steel beads (Qiagen) and 150 μL DMEM supplemented with 2% FBS and homogenized using a TissueLyser II (2 cycles at 30 Hz, 1 min, Qiagen). Homogenates were clarified by centrifugation and frozen until plaque assay titration. Viral loads in the saliva were further amplified on BHK-21 or C6/36 for 3 days, and presence of the virus was determined by direct visualization of cytopathic effects or RT-qPCR detection in the conditions described above for competition assays.

### Human blood and ethics statement

Human blood used to feed mosquitoes was obtained from healthy volunteer donors. Healthy donor recruitment was organized by the local investigator assessment using medical history, laboratory results, and clinical examinations. Biological samples were supplied through participation of healthy volunteers at the ICAReB biobanking platform (BB-0033-00062/ICAReB platform/Institut Pasteur, Paris/BBMRI AO203/[BIORESOURCE]) of the Institut Pasteur to the CoSImmGen and Diagmicoll protocols, which have been approved by the French Ethical Committee (CPP) Ile-de-France I. The Diagmicoll protocol was declared to the French Research Ministry under the reference: DC 2008-68 COL 1. This study was carried out in strict accordance with the recommendations in the Guide for the Care and Use of Laboratory Animals of the National Institutes of Health. The research protocol was approved by the Institutional Animal Care and Use Committee (IACUC) of Virginia Tech.

### *In silico* N-glycosylation predictions

N-glycosylation prediction was performed using two webservers: NetNGlyc 1.0 (http://www.cbs.dtu.dk/services/NetNGlyc/) and NGlycPred (https://bioinformatics.niaid.nih.gov/nglycpred/). For both servers, we used the MAYV E2 glycoprotein sequence from strain IQT4235. With the NetNGlyc server, we used the default option to run predictions only on the Asn-Xaa-Ser/Thr sequons and used a 0.5 threshold. In NGlycPred server, we used the preferred option to consider structural properties and patterns.

### Molecular dynamics simulations and calculation of binding free energy values

All the structural models of MAYV in complex with the human Mxra8 receptor were built based on the coordinates position of CHIKV crystallographic structure in complex with Mxra8 (PDB ID: 6JO8 [46]). To model the complexes between MAYV E2 subdomain and Mxra8 D2 domain, we removed from 7KO8 and 6JO8 structures all residues not included in those domains. The asymmetric unit of MAYV Cryo-EM structure (PDB ID: 7KO8) has 4 E1-E2 heterodimer copies whereas CHIKV crystallographic structure (PDB ID: 6JO8) has three copies. Thus, we cross-aligned, one by one, the E2 B subdomains of MAYV structure to each E2 B subdomains of CHIKV with the Chimera software [67]. This cross-alignment returned 12 complexes between MAYV E2 B subdomains and Mxra8 D2 domain. We estimated predicted free energy changes upon T179N mutation for all 12 MAYV complexes using FoldX 5 software [68]. Before energy calculations, each structure was treated to optimize sidechain rotamers using the FoldX function *RepairPDB*. In FoldX, wild-type structures were mutated and the ΔΔG (in kcal/mol) was estimated by subtracting the resulting energy of the mutated complex by the wild-type complex. We reported the FoldX result as mean ± standard deviation of ΔΔG for the 12 MAYV complexes. These same WT and T179N complexes were used as initial structures for molecular dynamics simulations using Amber 20 suit of programs with the *ff14SB* force field. Using *tleap*, we added Na+ and Cl-counter ions to reach net-neutralization with salt excess to reach 150 mM NaCl [69]. Each structure was solvated in a truncated octahedral box (15 A from the solute) filled with *TIP3P* water. The *PBRadii* was set to *mbondi2* with *tleap* program. The system was minimized by 2500 steps of steepest descent minimization and 2500 steps of conjugate gradient. The equilibration was performed using the NVT ensemble (200 ps) followed by NPT ensemble (200 ps), both with harmonic restraints in protein atoms. A last NPT equilibration step without restraints was performed for 500 ps. For each complex, the production run was performed at 298 K for 2 ns with a time step of 0.002 fs in triplicate. Hydrogen-containing bonds were constrained using SHAKE [70]. Long-range electrostatic interactions were calculated using particle Mesh Ewald and short-range nonbonded interactions were calculated with a 9 A cutoff. In all simulation steps we applied a harmonic restraint of 10 kcal·mol-1 ·Å −2 to the backbone atoms to prevent an overall displacement of the complex from its initial position. We estimated the relative binding free energy using 1-trajectory Molecular Mechanics Generalized Born Surface Area (MM/GBSA) calculations. For this, we used 20 frames from entire 2 ns simulations with MMPBSA.py script [71]. The individual topology files for the calculations were generated using ante-MMPBSA.py script. Since our objective was only to compare wild-type and mutant interactions, we did not include the entropy term. As result, the binding free energy estimate reported a relative value, not an absolute. In MMGBSA calculations, we used igb model 5 (all the other parameters were default). We reported the MMGBSA results as a ΔΔG of the difference between the relative ΔG estimated for T179N MAYV:Mxra8 and WT MAYV:Mxra8. To model the entire MAYV spike, formed by three E1-E2 heterodimers (not including transmembrane domains), in complex with Mxra8, we aligned individually the MAYV E1-E2 heterodimers (from PDB ID: 7KO8) with CHIKV E1-E2 heterodimers (from PDB ID: 6JO8) in Chimera. Then, we created the MAYV T179N mutated structure with the YASARA software [72]. Both wild-type and mutant MAYV trimeric complexes were submitted to energy minimization in solvent with YASARA to obtain an initial model. Then, we performed a 20 ns molecular dynamics simulation in quintuplicate for WT and T179N MAYV complexes, following the same simulation protocol described above without backbone restraints. The MMGBSA calculation was performed using 100 frames from the last 4 ns with the same parameters described above. Before any calculations or simulations, the protonation state at pH 7.4 for each residue of the structural models was assigned using the *pdb2pqr30* script [73] and *propka* as the titration state method [74]. For all cases, we also modeled the missing loop of Mxra8 D2 domain using YASARA software [72]. The convergence of the simulations were accessed by the RMSD of the Mxra8 backbone in a trajectory aligned by the MAYV backbone (**Fig. S7**).

**Figure S7.**
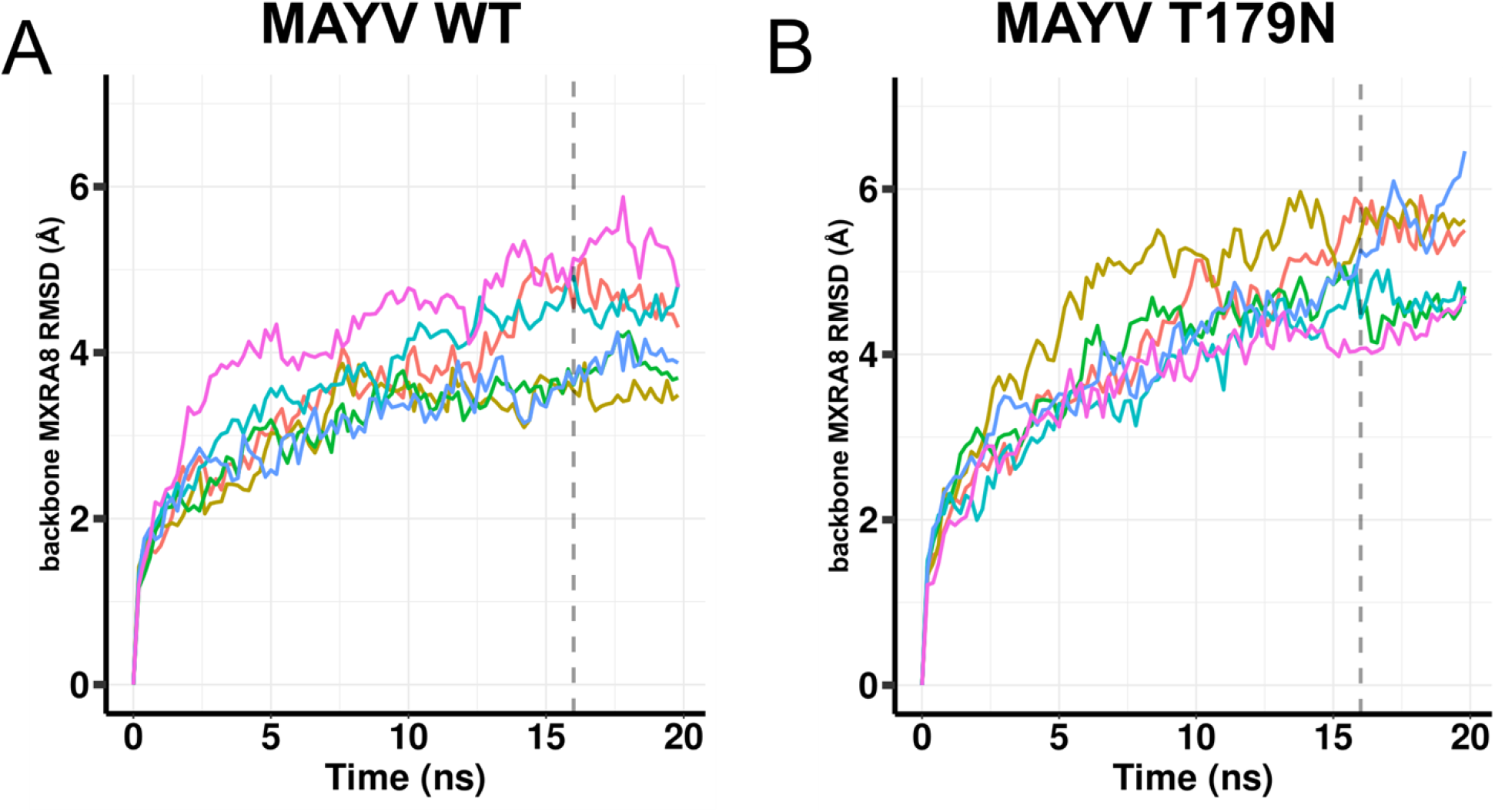
Root-mean-square deviation (RMSD) of Mxra8 receptor in complex with (A) MAYV WT spike or (B) MAYV T179N during 20 ns molecular dynamics simulations. To calculate the RMSD of Mxra8 backbone atoms, the trajectory was aligned by the MAYV backbone atoms using the first frame as reference in VMD. Each line corresponds to a replicate and the vertical dashed line indicates the last 4 ns of the simulation.

### Statistical analysis

All data were analyzed using the Prism 9 software (GraphPad) and are presented as mean ± standard deviation unless indicated otherwise. The statistical tests used are described in each figure legend and were performed using GraphPad Prism. All experiments (except serial and DMS passaging) were performed in two to three independent replicates with at least three technical replicates per group.

## Acknowledgements

The authors thank Rodrigo Guabiraba for helpful discussions. Illustrative figures in this manuscript were created with BioRender.com. This work was funded by the DARPA PREEMPT program managed by Dr. Rohit Chitale and Dr. Kerri Dugan and administered through DARPA Cooperative Agreement HR001118S0017 (the content of the information does not necessarily reflect the position or the policy of the U.S. government, and no official endorsement should be inferred). This work also received funding from Laboratoire d’Excellence Integrative Biology of Emerging Infectious Diseases (grant ANR-10-LABX-62-IBEID) to M-C.S. and M.V. Further support was provided by startup funds awarded to J.W-L by the Virginia-Maryland College of Veterinary Medicine and a grant from the One Health Research Funding Program awarded to J.W-L and P.M.

